# Seasonal drought timing shapes flowering phenology directly and through biotic interactions

**DOI:** 10.1101/2025.07.16.665144

**Authors:** Barel Tsafon, Or Gross, Niv DeMalach

**Affiliations:** Institute of Plant Sciences and Genetics in Agriculture, Faculty of Agriculture Food and Environment, The Hebrew University of Jerusalem, Rehovot, Israel

**Keywords:** Phenology, Climate change, Niche partitioning, Competition, Seasonality, Flowering, Drought timing, Phenotypic plasticity, Mediterranean ecosystems, Annual plants

## Abstract

Flowering time underpins plant fitness, species coexistence, and ecosystem functioning. While global warming consistently advances flowering, the influence of water availability remains unclear. We hypothesized that this inconsistency reflects the overlooked timing of drought. In 200 experimental Mediterranean annual-plant communities, we imposed early-, mid-, and late-season dry periods and grew plants in monocultures and mixtures to disentangle physiological and competition-mediated responses. Early and late droughts shortened flowering duration: early drought delayed onset, late drought advanced termination. Some shifts were direct, others emerged through competition. A new community-level index revealed greater phenological segregation in mixtures, showing that plasticity alone can generate niche separation under competition. Both early and late droughts further enhanced this segregation. Together, our results demonstrate that the seasonal timing of drought governs flowering responses through both direct physiological pathways and indirect biotic interactions, emphasizing rainfall seasonality as a key driver of ecological responses to climate change.

**Impact statement:** Since flowering time is crucial to ecosystem functioning, shifts in the timing of drought could have far-reaching effects on the performance and resilience of ecological communities. Our study shows that droughts’ effect on flowering is dependent on their timing and influenced by interactions with neighboring species. This means that understanding how plants respond to changing water conditions requires looking beyond single species, considering both seasonality and community interactions.

## Introduction

Flowering phenology, the timing of flowering onset and termination, is a fundamental biological trait that shapes individual fitness and underpins key ecosystem functions and services (Kantsa et al., 2018; Taylor et al., 2019; Ren et al., 2024). The timing of major flowering events has broad public relevance with various cultural and economic consequences globally (Winkler and Brooks, 2020; Graves et al., 2017). Moreover, since flowering time influences competition for pollinators and other resources, it is recognized as a critical component of niche partitioning, where temporal segregation across the growing season reduces competitive overlap and promotes species coexistence (Sherry et al., 2007).

There is mounting evidence that climate change is altering phenological patterns across ecosystems, with cascading consequences for community structure, plant–pollinator interactions, and ecosystem stability (Liu et al., 2025; Fitter and Fitter, 2002; Vitasse et al., 2022; Høye et al., 2013). Hence, flowering phenology is often viewed as a direct indicator of ecosystem change (Alexander and Levine, 2019; Cleland and Wolkovich, 2024).

It is well established that increased temperature triggers earlier flowering phenology (Cook et al., 2012; Peñuelas and Filella, 2001; Wolkovich et al., 2012; Williamson et al., 2025), but the effects of changes in precipitation patterns are unclear, with conflicting results (Knapp et al., 2024; Lu et al., 2023; Martén-Rodríguez et al., 2025; Matthews and Mazer, 2016). In seasonally dynamic ecosystems, such as Mediterranean systems, water availability is a limiting factor, and the impact of drought may depend strongly on its timing within the growing season (Ru et al., 2024; Gordo and Sanz, 2010; Johansson et al., 2013; Reyer et al., 2013). It is now clear that climate change is causing these ecosystems to experience a shortening of the wet season, along with shifts in rainfall timing and distribution (Delworth et al., 2022; Vicente-Serrano et al., 2025). However, so far, experimental studies of flowering phenology have focused on altering the total rainfall amount rather than its timing (Castillioni et al., 2022; Van Dyke and Kraft, 2025).

The impact of climate change on plants is recognized as both direct, through physical changes, and indirect, through biotic interactions (Chu et al., 2016). Unlike temperature, the water available to a plant is influenced not only by abiotic factors but also through depletion caused by neighboring individuals (Trautz et al., 2017). This makes interspecific competition an even more critical consideration, yet one that very few phenology experiments have addressed. The importance of this perspective is particularly evident in the case of flowering, with empirical work showing that temporal patterns shift as species diversity changes (Wolf et al., 2017). Ultimately, two key questions arise: first, how do plants adjust their flowering phenology in response to temporal changes in water availability, and second, can competitive interactions between species modify this effect?

Here, we experimentally manipulated the timing of water availability in controlled settings to test its effects on species-level flowering phenology and community-level phenological overlap (Fig. 1). We established annual plant communities subjected to a control and three drought treatments (early-season, mid-season, and late-season), each involving the same reduction in total water supply, thereby mimicking the shortening of the rainy season observed in Mediterranean systems worldwide (Vicente-Serrano et al., 2025). Each species was grown in monocultures and mixed communities to disentangle direct physiological responses to treatments from effects mediated by interspecific interactions (hereafter, ‘competition’). In addition, we developed a new index to capture the degree of multi-species niche partitioning at the community level.

**Figure 1.**
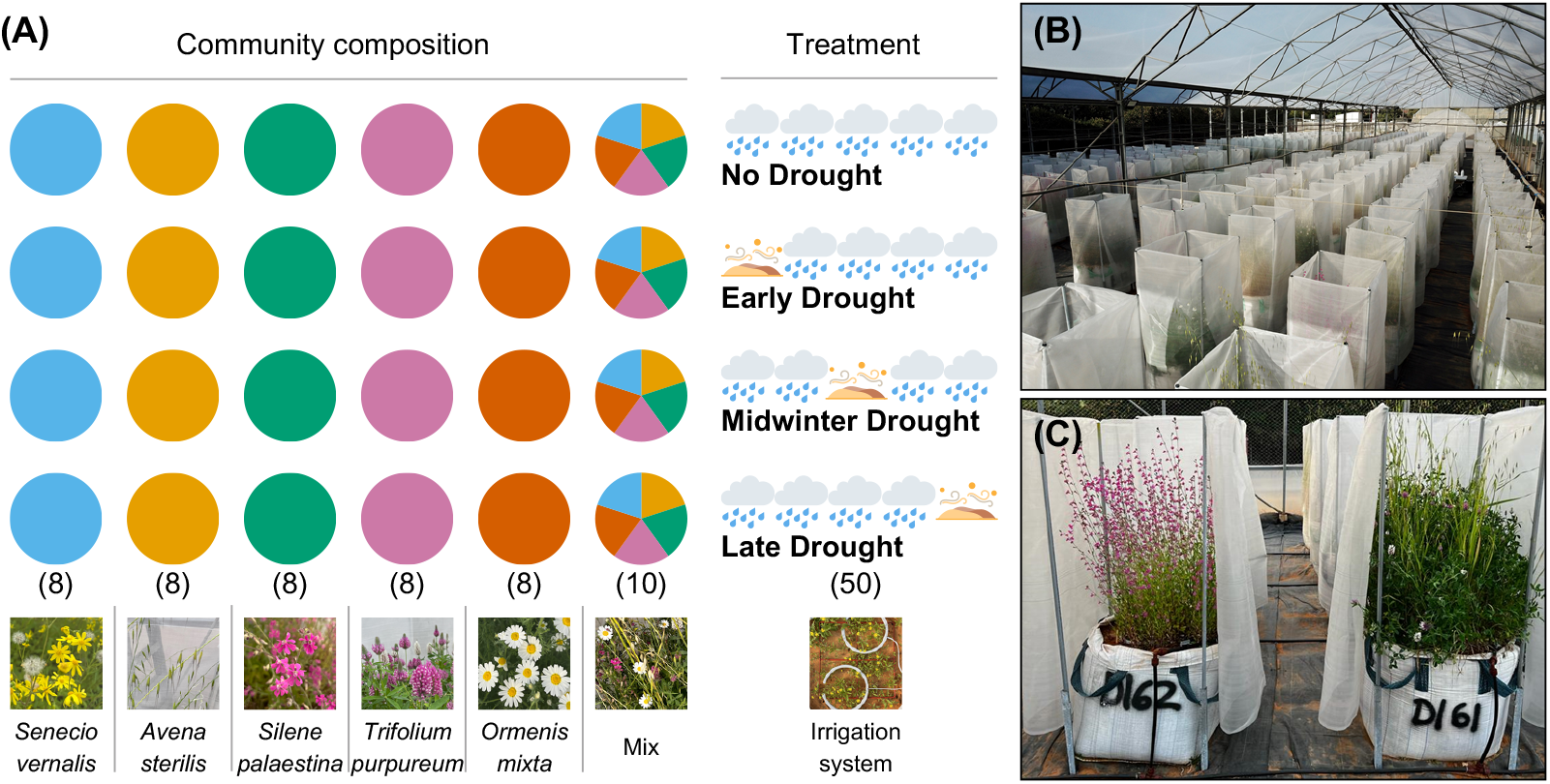
Overview of the experimental design. (A) Schematic illustration of the experimental treatments. Each circle represents one community; colored circles correspond to monocultures of five focal species (eight replicates each), and the multicolored pie chart represents mixed species communities (ten replicates). Drought treatments include a no drought (control), early season drought, mid-season drought, and late season drought. (B) Photograph of the greenhouse showing the full set of experimental units. (C) Example of two experimental communities: a *Silene* monoculture (left) and a five-species mixture (right).

Four hypotheses were tested: (1) Drought effects are both direct and mediated by species interactions; (2) Early-season drought delays flowering onset due to postponed germination, thereby reducing the flowering period; (3) Late-season drought causes early ending of flowering (hereafter offset) due to early-season termination, also reducing the flowering period; (4) Early- and late-season droughts constrain phenological extremes, thereby reducing phenological niche partitioning and increasing overlap.

## Materials and methods

### Establishment of the experiment

The experiment started in October 2022 in a greenhouse located at the agricultural campus of the Hebrew University of Jerusalem in Rehovot, Israel (31.90^*◦*^, 34.80^*◦*^). The greenhouse had a polyethylene roof to prevent natural precipitation and net walls (20 cm *×* 20 cm mesh opening) that enabled maximum ventilation with temperatures close to ambient. The climate in the region is Mediterranean with mild rainy winter (*∼*560 mm y^*−*1^, falling mainly between October and April) and a dry, hot summer (Issaka et al., 2023).

The experimental communities mimicked the natural annual plant communities growing on the red sandy soil of Israel’s coastal region (Hamra communities, (Issaka et al., 2023)). We chose five common species varying in phenology, functional group, and taxonomic group: (1) *Senecio vernalis*, (Compositae), typically flowering from December to February. (2) *Avena sterilis*, (Poaceae), flowering from March to May; (3) *Silene palaestina*, (Caryophyllaceae), flowering from March to April; (4) *Trifolium purpureum* (Fabaceae), flowering from March to April; (5) *Ormenis mixta*, (Compositae), flowering from April to May. All seeds were collected in the spring of 2021 from nearby natural sites.

The experiment included 200 units, communities grown in industrial bags (0.7 m*×*0.7 m*×*0.7 m) spaced at least 0.8 m apart. Every bag was surrounded by a 50-mesh net that rose one meter above the soil surface to minimize dispersal. Hamra soil was collected from a depth of at least one meter in the area surrounding the experimental site, to minimize the presence of a natural seed bank and ensure that only sown species emerged. It was fertilized with 5.26 g m^−2^ Osmocote Pro fertilizer (2 g N m^−2^) to compensate for the lack of organic matter at that depth of this nutrient-poor soil.

Each species was grown in a monoculture or together with the other four species in mixture communities (Fig. 1). Monocultures enable the quantification of the direct impact of treatments on any of the species, while the difference between mixtures and monocultures allows the estimation of the competition-mediated effects. Based on the preliminary germination trials, we sowed seeds aiming for an initial density of 1000 seedlings per bag (in mixture units: 200 seedlings per species).

On October 10th, we sowed the seeds of most treatments just before the start of the irrigation schedule. In the early drought treatment that aimed to germinate later, we sowed the seeds on November 16th to avoid seed loss before irrigation.

### Experimental schedule

To ensure that each community received a separate treatment, irrigation was applied through the NetBow™ drip irrigation system (by Netafim Ltd.). Two NetBow™ rings were installed in each bag (flow rate was 16 mm/h per bag). Irrigation events occurred once a week to mimic the natural cycles of wetting and drying. Each event lasted 1.5 h; thus, every week, the communities received 24 mm of water: 576 mm throughout the long season (control) and 456 mm in the drought treatments. Additionally, during the first three weeks after sowing, water was lightly added with a hose to enhance germination.

We created an irrigation schedule based on natural precipitation in the region, which historically lasts (on average) between mid-October and the beginning of April. However, in the last 30 years, the number of rainy days has decreased by 10% (Drori et al., 2021). Therefore, we chose to examine a 20% reduction compared to the historical record. We designed the experiment such that all drought treatments would have the same number of irrigation events and amount of water, to ensure that changes in phenology can be causally attributed to the timing of water stress within the growing season.

Each experimental unit was assigned to one of four drought types: (1) “no drought”: an irrigation period of 25 weeks (control); (2) “early drought”: a shorter irrigation period of 20 weeks, starting five weeks after the control representing years where the growing season is delayed due to lack of rainfall in Autumn; (3) “midwinter drought”: a normal irrigation period of 25 weeks with a five-week drought in January; (4) “late drought”: an irrigation period of 20 weeks, starting as the control and ending five weeks before the control. Experimental units were randomly assigned to create eight monoculture replicates for each species per treatment and ten replicates of mixtures (Fig. 1).

At the beginning of the first and last months of irrigation, the number of plants was estimated inside every bag. Counts were made in 40 cm *×* 40 cm quadrats at the center of each bag. Flowering phenology was tracked in each unit once a week in a binary survey throughout the season. Units were considered flowering if at least 1% of individuals showed open flowers. The 1% threshold was calculated using the measured densities and sampled across the entire surface area of the bag. During drought treatments, soil moisture was measured weekly in every bag with a soil moisture meter to assess the treatments’ effect on soil water content (Fig. S6). Measurements were taken from a depth of 10 cm, since many annuals have a shallow root system. We measured at fixed locations in the corners of the quadrats to avoid edge effects while keeping the integrity of the communities’ center.

### Statistical analysis

All analyses were conducted in R version 4.5.0. We used linear regression to test the effects of drought treatments (coded using dummy variables), interspecific competition (monoculture vs. mixture), and their interactions on several response variables. These included flowering onset, flowering offset, and length of the flowering period. To quantify community-level niche partitioning of the flowering season, we used the weekly flowering observations described above. For each community, we defined the flowering season as the number of weeks between the onset of the first flowering species and the offset of the last. We then calculated a standardized niche partitioning index for each community. The index ranges from 0 (no partitioning, complete overlap) to 1 (maximum partitioning, no overlap), and is defined by the following equation:

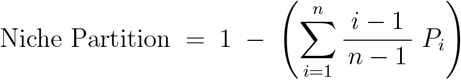

where *n* is the number of species in the community, *i* is the number of flowering species, and *P*_*i*_ is the proportion of the flowering season during which *i* species flower. The index assigns greater weight to weeks with more co-flowering species, capturing the degree of phenological overlap. Subtracting this overlap score from one yields a measure of temporal niche partitioning at the community level.

The niche partitioning index was calculated per unit for the real mixed communities, reflecting the co-flowering of species grown together. In contrast, monoculture communities were composed of single species grown in isolation, so we constructed simulated mixed communities by randomly selecting one monoculture replicate per species and combining their flowering data to mimic the structure of a five-species mixture, but without interspecific competition (Fig. 3C). While many such combinations are possible, most would not be statistically independent. To address this, we constructed eight non-overlapping simulated mixed communities in each iteration, ensuring that each monoculture replicate appeared only once. This preserved independence across communities while maintaining the original sample size.

We ran a linear regression to estimate treatment effects, including the real and simulated mixed communities in each iteration. The procedure was repeated 100 times to minimize the stochastic effects of sampling from the monocultures. Parameter estimates from these iterations were averaged to robustly estimate treatment effects. However, since *p*-values cannot be simply averaged, we applied the Cauchy combination test (CCT) using the STAAR R package to compute integrated *p*-values across iterations.

## Results

The phenological order of species in the experiment matched their natural flowering sequence. In all treatments, *Senecio* was the first to flower, with flowering onset within five weeks of irrigation initiation, followed by *Avena, Silene, Trifolium*, and finally *Ormenis*, which continued to flower well beyond the end of irrigation (Fig. 2A). Interspecific competition shortened the flowering period of *Senecio, Silene*, and *Avena* by 3 to 8 weeks, as revealed by comparisons between their phenology in monocultures and mixtures (Figs. 2A, S1; Table S1). Regarding the effects of drought timing, both early- and late-season droughts shortened the flowering period, whereas mid-season drought had a negligible impact (Figs. S1; Table S1). Early drought, characterized by a five-week delay in irrigation, resulted in a species-specific response of 2- to 5-week delay in the onset of flowering across all species (Fig. S2; Table S2). However, as this treatment did not affect offset (except in *Avena* monocultures), it led to a shorter overall flowering period (from the onset of the earlier species to the offset of the latest species, Figs. S1, S3). Late drought caused early offset in the mid-season species, *Avena* and *Silene*. In *Avena*, this effect was consistent across monocultures and mixtures, suggesting a direct response to the treatment (Fig. 3; Table S3). In contrast, for *Silene*, the effect was limited to mixtures, implying only an indirect impact of the treatment, mediated by interactions with other species (Fig. 3; Table S3).

**Figure 2.**
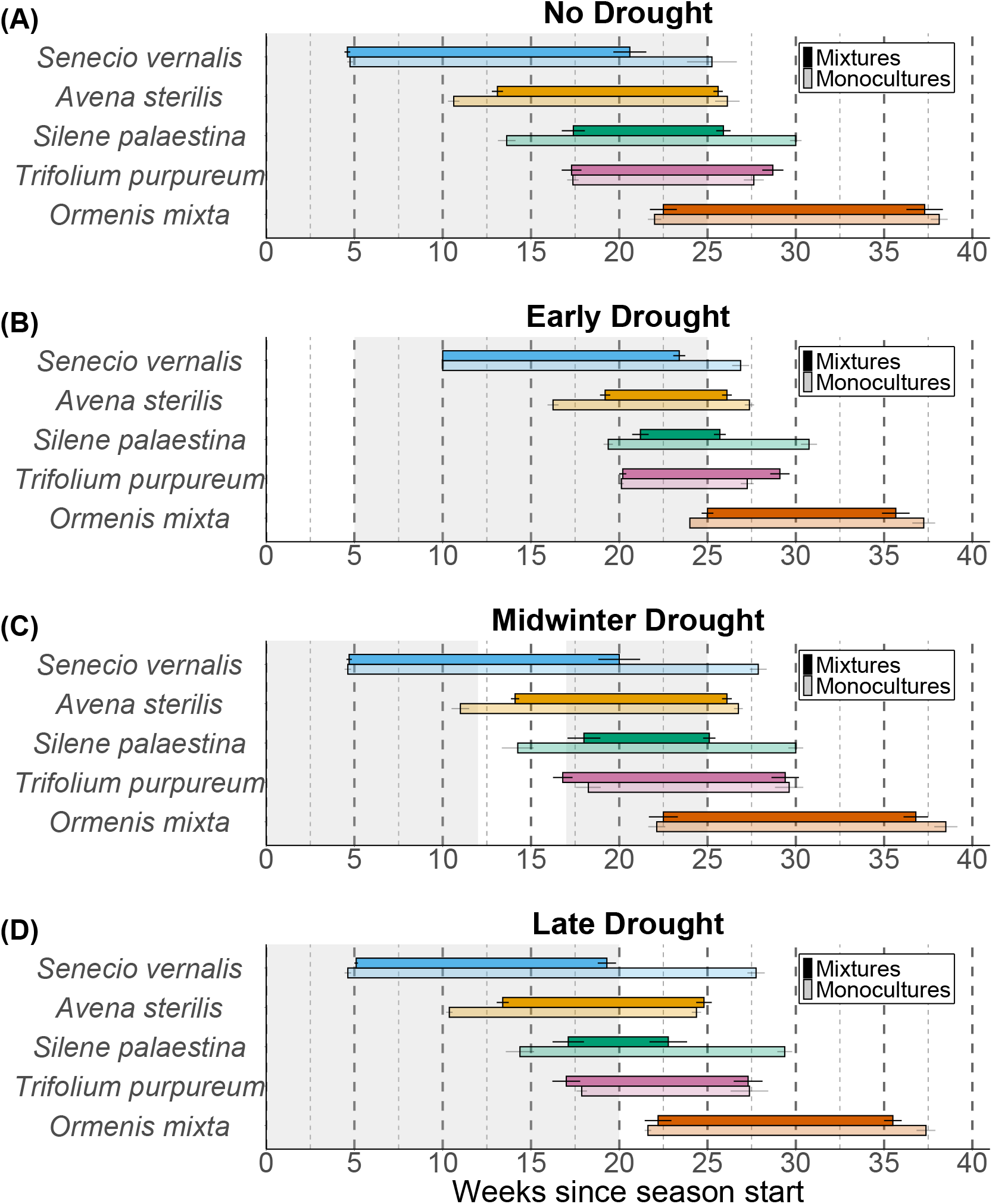
Drought and competition alter flowering phenology. Calendric flowering phenology of each species in monocultures and mixtures under the four drought treatments (A)–(D). Each color represents one species. Light and dark shades indicate monocultures and mixtures, respectively. The x-axis shows the number of weeks since the start of the season in mid October (week 0 marks the onset of irrigation). Gray areas indicate periods with irrigation, while white areas indicate drought. Error bars represent standard errors of the mean.

**Figure 3.**
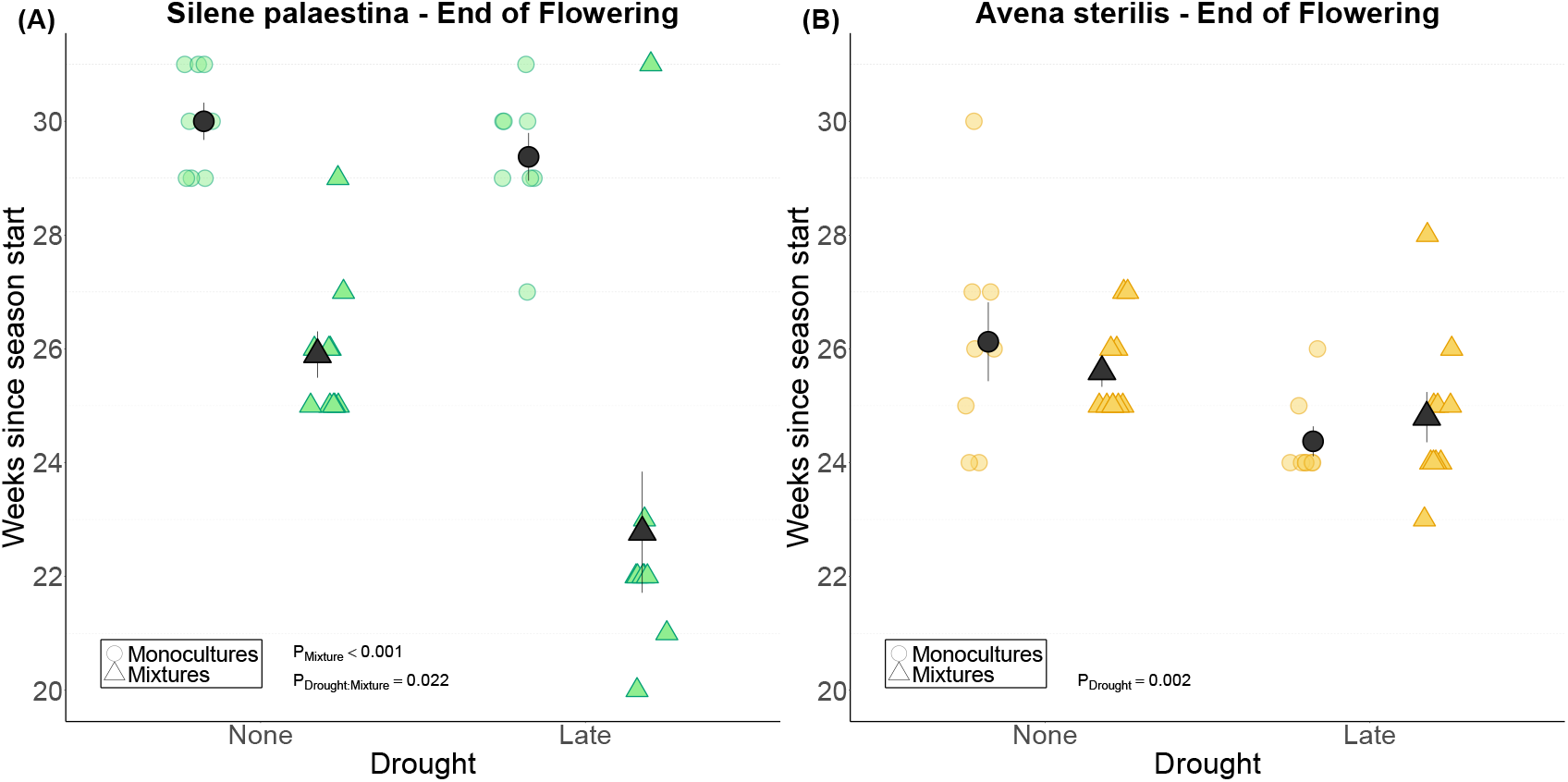
Direct and indirect effects of drought on the ending of the flowering period for (A) *Silene palaestina* and (B) *Avena sterilis*. The x-axis indicates the watering treatments. Circles represent monocultures, and triangles represent mixtures. Error bars represent standard errors of the mean.

Interspecific competition increased the fraction of the flowering season with only one species flowering from 40% to 51%, thereby reducing the proportion of multispecies overlap (Fig. S4). This was accompanied by a sharp decline in the proportion of time when all species flowered simultaneously, from 11% in monocultures to 3% in mixtures. The shift toward lower overlap was driven by shortened flowering durations in species that typically flower early or mid-season, while the overall length of the flowering season remained unchanged (Fig. 2A). Early-season drought further reduced the proportion of time during which multiple species flowered simultaneously (Fig. S4). This was primarily due to a delayed onset in mid-season species, while their offset remained unchanged (Figs. 2B, S2, S3; Tables S2, S3). As a result, their overlap with the late-flowering *Ormenis*, which also exhibited a delayed onset, was reduced. Finally, the late-season drought reduced the number of days when four or five species flowered simultaneously (Fig. S4), as mid-season species, particularly *Silene* and *Avena*, terminated flowering earlier than under control conditions (Fig. 3).

To quantify phenological niche partitioning at the community level, we developed a new index (Fig. 4A; see Methods for details). Existing approaches typically assess overlap between species pairs, but we were not aware of indices that capture temporal overlap across entire communities. Our index is based on the number of species flowering each week throughout the season (Fig. S4). Weeks with more co-flowering species indicate higher overlap, while weeks with fewer species indicate greater temporal segregation. The index is standardized between zero (maximum overlap, all species flower simultaneously) and one (maximum partitioning, each species flowers at a different time).

We compared niche partitioning in mixtures to expected values derived from monocultures. By randomly assembling the phenology data of monocultures into simulated mixed communities, we constructed a null model that excludes competition and enables the detection of direct drought effects (Fig. 4A; see Methods for details). The niche partitioning was higher in the real mixtures compared to the simulated mixtures by *∼*0.1, equivalent to a 10-percentage point increase, across all treatments, indicating that interspecific competition enhanced the niche partitioning of flowering periods. Both early and late drought further intensified niche partitioning in the real mixed communities, increasing the index by *∼*0.04 (Fig. 4B; Table S4). In the monoculture-based simulated mixed communities, the effect of early drought was similar to that observed in the real mixtures, with an estimated interaction very close to zero (Table S4), suggesting a primarily direct effect. In contrast, under late drought, the increase in niche partitioning was *∼*0.04 smaller in the simulated mixtures than in the real ones, suggesting a potential indirect effect of competition, although this interaction was not statistically significant (*P* = 0.14), likely due to insufficient statistical power. Calculating the community niche partitioning using the classical approach based on mean pair-wise partitioning yielded similar results: the index of real mixtures was *∼*0.1 higher than simulated mixtures, and late drought further increased it (Fig. S5).

**Figure 4.**
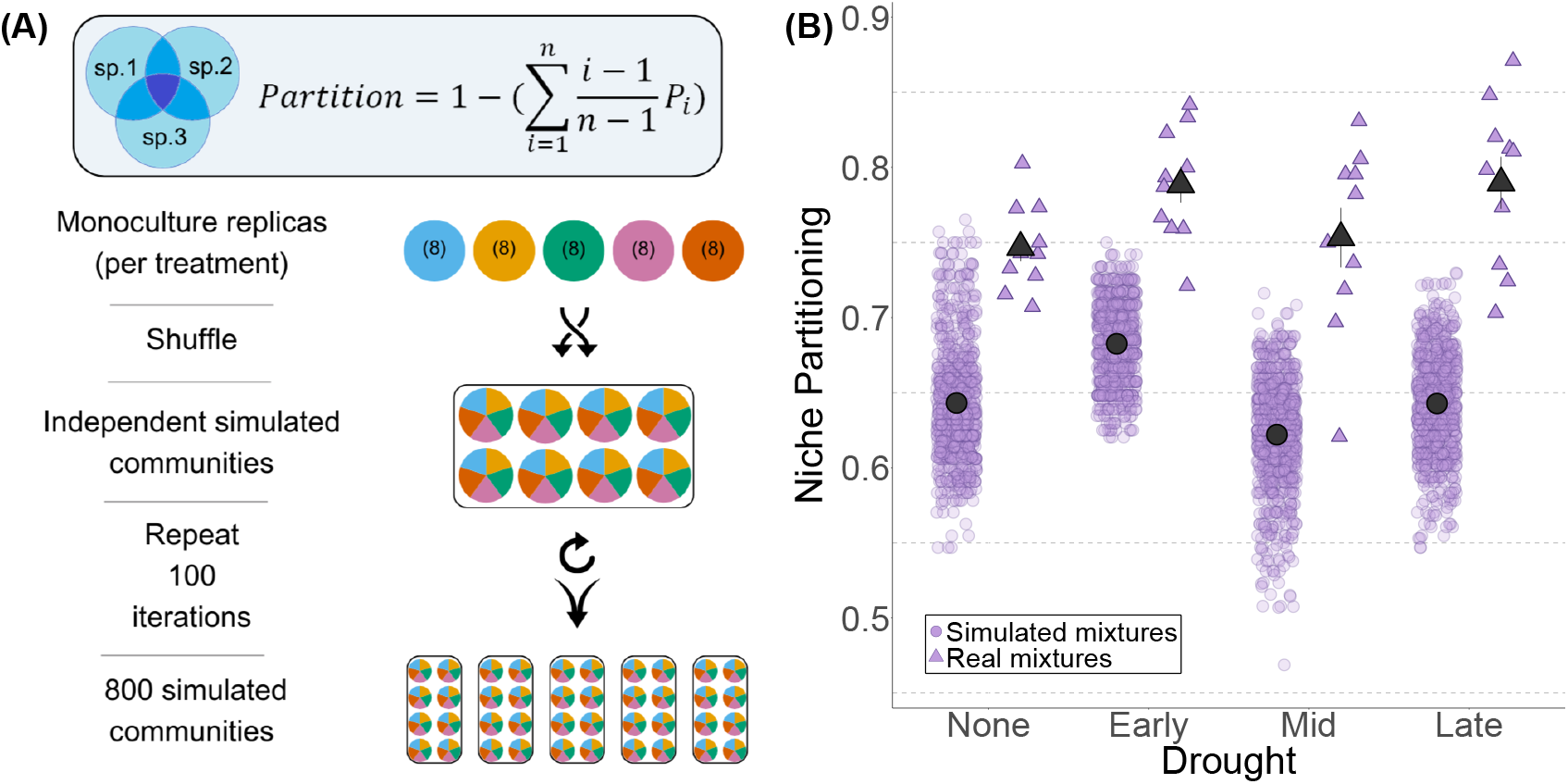
Drought and competition increase phenological niche partitioning. (A) Overview of the niche partitioning index. The panel includes the equation used to calculate the index (see Methods), and a schematic representation of the simulation process for one treatment. (B) Effects of drought treatments on the niche partitioning index. Triangles are mixed communities, and circles are simulated mixed communities calculated based on monocultures. Error bars show standard errors of the mean. pEarly = 0.033, pLate = 0.028, pMono < 0.0001.

## Discussion

We found that the timing of drought shapes flowering phenology at the species level and phenological niche partitioning at the community level, with these effects arising through two mechanisms: directly in a species-specific manner and indirectly through interspecific competition. Early-season drought delayed the flowering onset of early- and mid-season species, and reduced their overlap with late-season species. In contrast, mid-season drought had no detectable effect on the onset or termination of flowering. Late-season drought led to an earlier end of flowering, both directly (as seen in *Avena*) and indirectly through altered competitive dynamics (as in *Silene*), resulting in increased phenological niche partitioning at the community level. Our findings also highlight the often-overlooked role of competition-induced phenotypic plasticity in shaping flowering phenology. Competition consistently shortened the flowering duration of most species, which contributed to increased phenological niche partitioning at the community level.

### Competition and phenological niche partitioning

In classical ecology, character displacement and limiting similarity describe an evolutionary process in which species diverge in their traits to reduce competition for shared resources (Macarthur and Levins, 1967). This process often results in fixed differences in phenology, as observed in our monoculture treatments under control conditions, where species exhibited distinct flowering periods. However, our findings show that phenological niche partitioning can also arise within a single growing season, through phenotypic plasticity in response to the immediate presence of other species. Indeed, previous work has demonstrated that plasticity, rather than adaptation, is the primary driver of flowering-time variation along climatic gradients, highlighting its capacity to mediate rapid phenological shifts (Ramirez-Parada et al., 2024). Our observation also aligns with recent theoretical work predicting that competition can induce plastic shifts in phenology (Levine et al., 2022; Rudolf, 2019).

Plasticity-driven niche partitioning has been well-documented for below-ground traits such as root allocation and nutrient uptake (Phoenix et al., 2020; Schiffers et al., 2011; Ashton et al., 2010), but it has rarely been studied in the context of flowering phenology. A notable exception is Wolf et al. (2017), who demonstrated that increased species richness led to greater variation in peak flowering time. This pattern was interpreted as evidence for niche partitioning driven by competition. Our results are consistent with their findings but differ in that we quantify partitioning based on the degree of overlap across the entire flowering period.

To date, two main approaches have been used to quantify phenological niche partitioning.

One focuses on the variance or distance among peak flowering times (Williams, 1995), while the other relies on the complement of the overlap in flowering duration, measured from onset to offset (Austin et al., 2024); both have advantages and disadvantages (Freitas and Bolmgren, 2008). Peak-based methods face several statistical challenges and often ignore important differences in the timing of onset and offset. Overlap-based methods are more informative but typically assess pairwise overlap between species. While intuitive, average pairwise overlap can obscure patterns that emerge at the community level.

To address these limitations, we developed a new index that quantifies the extent of temporal overlap at the community level. This approach captures simultaneous interactions among multiple species and provides a more holistic measure of phenological niche partitioning that fits better with our system. The close match between the community-level index and the mean pairwise metric in these simple communities (Fig. S5) validates our new measure and underscores its advantages as species richness and indirect interactions grow. Although designed for flowering phenology, the index is general and could be applied to any niche dimension where temporal or resource overlap occurs.

What drives plasticity-induced niche partitioning? One possibility is that plants sense neighboring individuals and adjust accordingly (Cahill et al., 2010). A simpler alternative is that plants respond to indirect environmental cues, such as declining water availability in mixtures (Wolf et al., 2017; Levine et al., 2024). The second explanation is supported by moisture patterns in our system (Fig. S6), which were consistently lower in mixtures than all monocultures except for *Trifolium*. In *Trifolium* monocultures, moisture levels were similar to those in mixtures, which may explain why it was unaffected by interspecific competition. Moreover, the phenological shifts observed under competition were consistent with each species’ typical flowering schedule. The early-season *Senecio* ceased flowering earlier under competition, the mid-season species *Avena* showed earlier offset, and *Silene* had both earlier onset and offset. In contrast, *Ormenis*, which flowers late and develops deeper roots, was unaffected by competition, consistent with theoretical predictions that late-season species are buffered from competition by rooting depth and reduced overlap (Levine et al., 2024).

### The Effects of Drought Timing

In accordance with our second hypothesis, early drought, achieved by delaying the onset of irrigation, resulted in later flowering onset across all species. This pattern suggests that a minimum plant size is required to trigger flowering (Wesselingh and de Jong, 1995). Since biomass accumulation depends on time, delayed germination leads to delayed flowering. However, plants partially compensated for the delay: a five-week postponement of irrigation resulted in a delay of less than five weeks in the onset of flowering for most species. This indicates that flowering is regulated by a combination of cues, not by soil moisture alone, strengthening prior knowledge (Andrés and Coupland, 2012).

We also hypothesized that late-season drought would lead to earlier flowering offset (hypothesis 3), and this prediction was supported for two species, *Silene* and *Avena*. Interestingly, early drought affected only flowering onset, while late drought affected only flowering offset. This indicates that the beginning and ending of the flowering period are not strictly linked, and that the total length of the flowering period is flexible rather than fixed (CaraDonna et al., 2014). In both cases, drought ultimately shortened the flowering period, in line with our expectations.

The early- and late-season droughts extended the dry season at the leading and trailing edges of the wet season, respectively, whereas the mid-season treatment imposed only a brief drought interval after plants had reached maturity. This mid-season pulse left both flowering onset and termination unchanged, indicating that established annuals can withstand shortlived water deficits without shifting their reproductive phenology (Castillioni et al., 2022; Fenollosa et al., 2024).

In contrast with our prediction that drought would reduce phenological niche partitioning (hypothesis 4), we found that it increased it. Our initial hypothesis was based on the assumption that early drought would primarily delay the flowering onset of early-flowering species, and that late drought would mainly trigger the early flowering offset of late-flowering species. Instead, we observed that early drought delayed the onset across all species, not just early bloomers, though the extent of the delay varied. Specifically, mid-season species showed a greater relative reduction in flowering duration. This led to an increased proportion of the flowering season dominated by single species, particularly early and late-season species, thereby enhancing phenological niche partitioning.

Similarly, the late drought treatment mainly affected mid-season species. This again reduced the periods of co-flowering and led to higher levels of niche partitioning. Notably, while the competitive species *Avena* (Issaka et al., 2023) showed a similar change of offset timing in both monocultures and mixtures, the drought-induced shift in *Silene* occurred only under competitive conditions, demonstrating the role of species interaction in mediating drought response and aligning with our first hypothesis.

## Conclusion

We have shown that early and late droughts significantly affect flowering time, while middle-season drought has little effect. This finding highlights the primary role of water availability timing in shaping phenology (Knapp et al., 2024). It is also relevant to the growing use of big data from herbarium records and citizen science in phenological research (Ahlstrand et al., 2025; Ramirez-Parada et al., 2024), as the timing of rainfall events may serve as a better predictor of ecological patterns than annual precipitation totals (Poppenwimer et al., 2023), especially in Mediterranean ecosystems where shifts in rainfall timing are more reflective of observed climate change (Delworth et al., 2022; Vicente-Serrano et al., 2025).

Lastly, our finding that drought effects are both direct and mediated by interspecific competition implies that responses cannot be fully understood from studies of single species. While the importance of indirect effects on community dynamics under climate change has been thoroughly recognized over the years (Adler et al., 2012; Van Dyke et al., 2022), phenology research has yet to adequately address these competition-mediated effects. Understanding how biotic interactions shape phenological responses is therefore essential for predicting the future structure and resilience of ecological communities in times of global climate change.

## Acknowledgements

The study was supported by the Israel Ministry of Environmental Protection (grant no. 22-1-18) and the Israeli Science Foundation (grants no. 2403/22 and 672/22) to N.D. Scholarships from the Jewish National Fund, the School of Environmental Studies of the Hebrew University, the Hebrew University Center for Sustainability, and the Women’s League for Israel supported B.T. and O. G. We thank Sara Barcza, Noa Levi, Shlomo Shik, Gal Ben Shlomo, Yuval Neumann, Yarden Schindler and Avishag Neeman for their assistance in fieldwork and Micha Mandel for statistical assistance.

