## Supplementary for "Seasonal drought timing shapes flowering phenology directly and through biotic interactions"

**Number of figures:** 6
**Number of tables:** 5

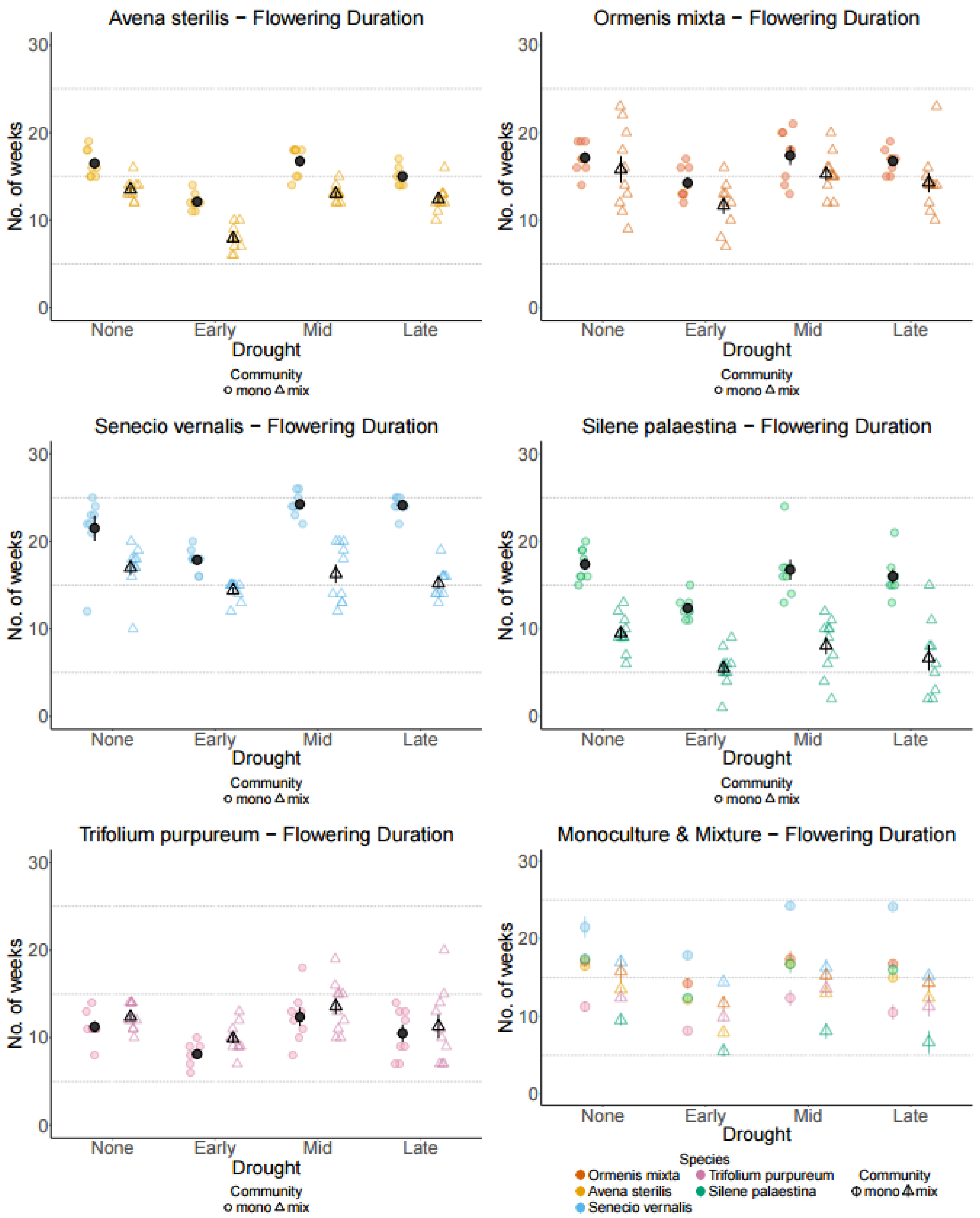


**Figure 1: The number of flowering weeks of all species under the four drought treatments (first year).** The y-axis is the number of weeks with open flowers. The x-axis is the treatments. Each species has its color. Round points are for the species in monocultures while triangular points are for the species in mixtures. The error bars are standard errors of means.

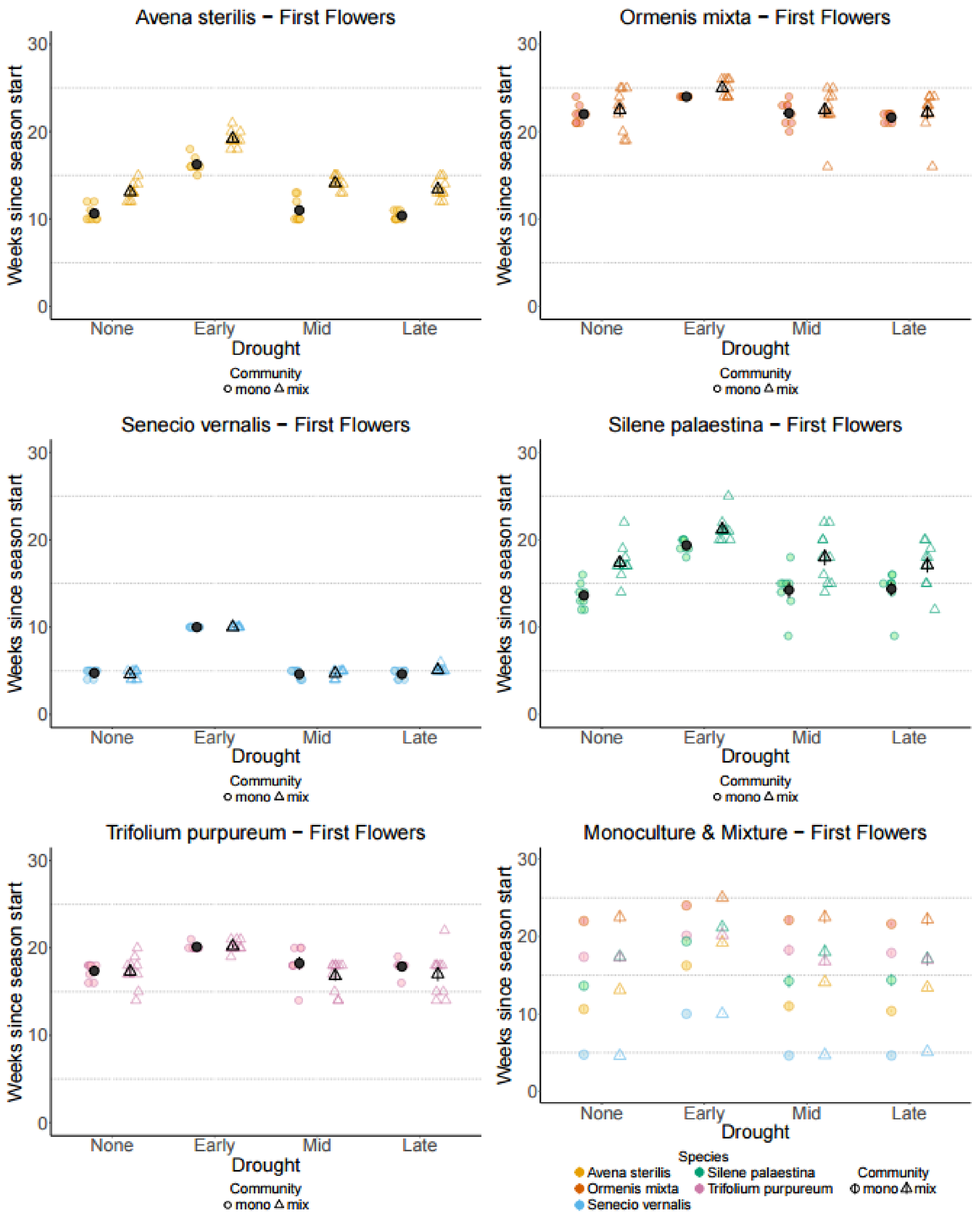


**Figure 2: The number of weeks from the season start until the onset of flowering of all species under the four drought treatments (first year).** The y-axis is the number of weeks since the first irrigation. The x-axis is the treatments. Each species has its own color. Round points are for the species in monocultures while triangular points are for the species in mixtures. The error bars are standard errors of means.

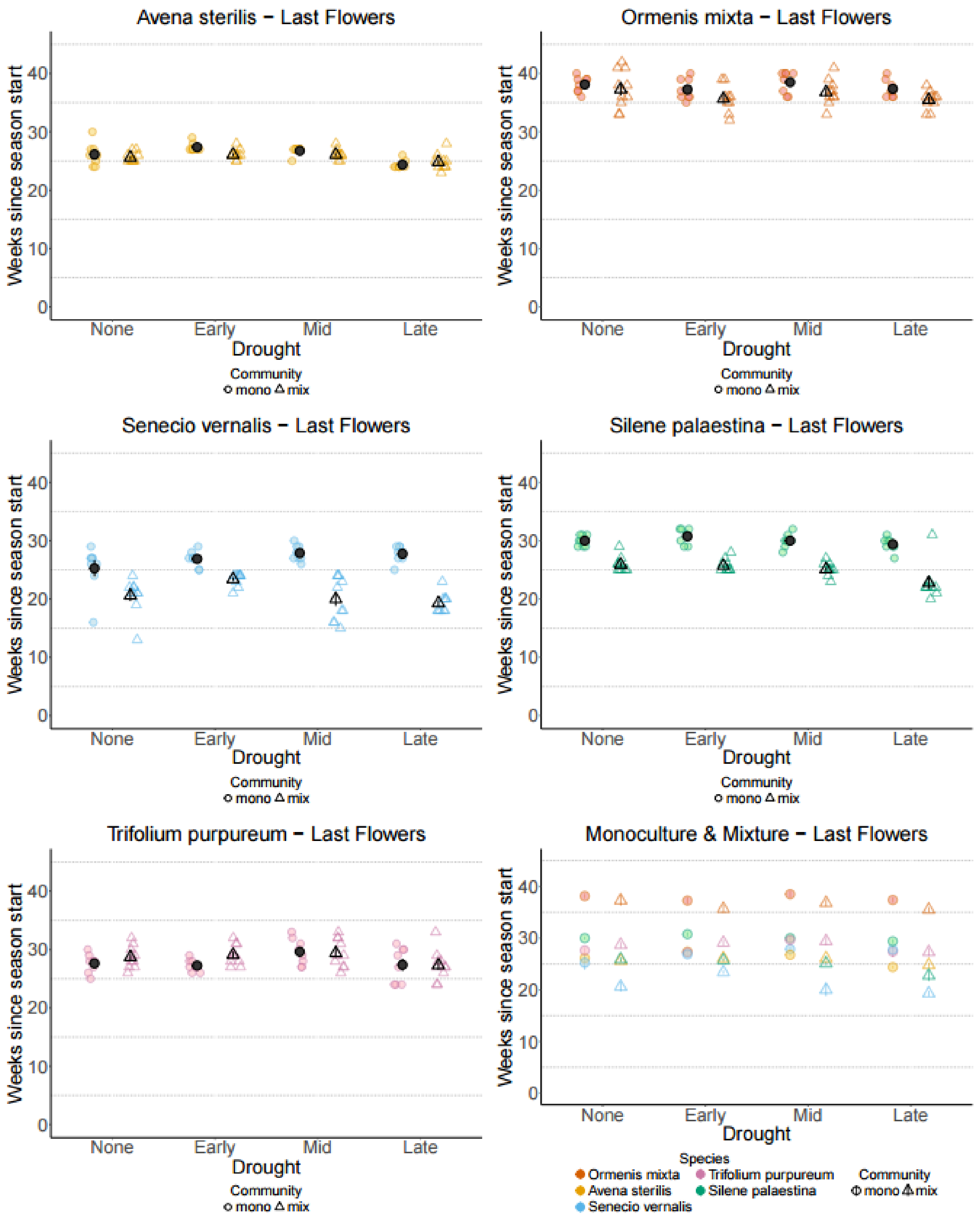


**Figure 3: The number of weeks from the season start until the end of flowering of all species under the four drought treatments (first year).** The y-axis is the number of weeks since the first irrigation. The x-axis is the treatments. Each species has its own color. Round points are for the species in monocultures while triangular points are for the species in mixtures. The error bars are standard errors of means.

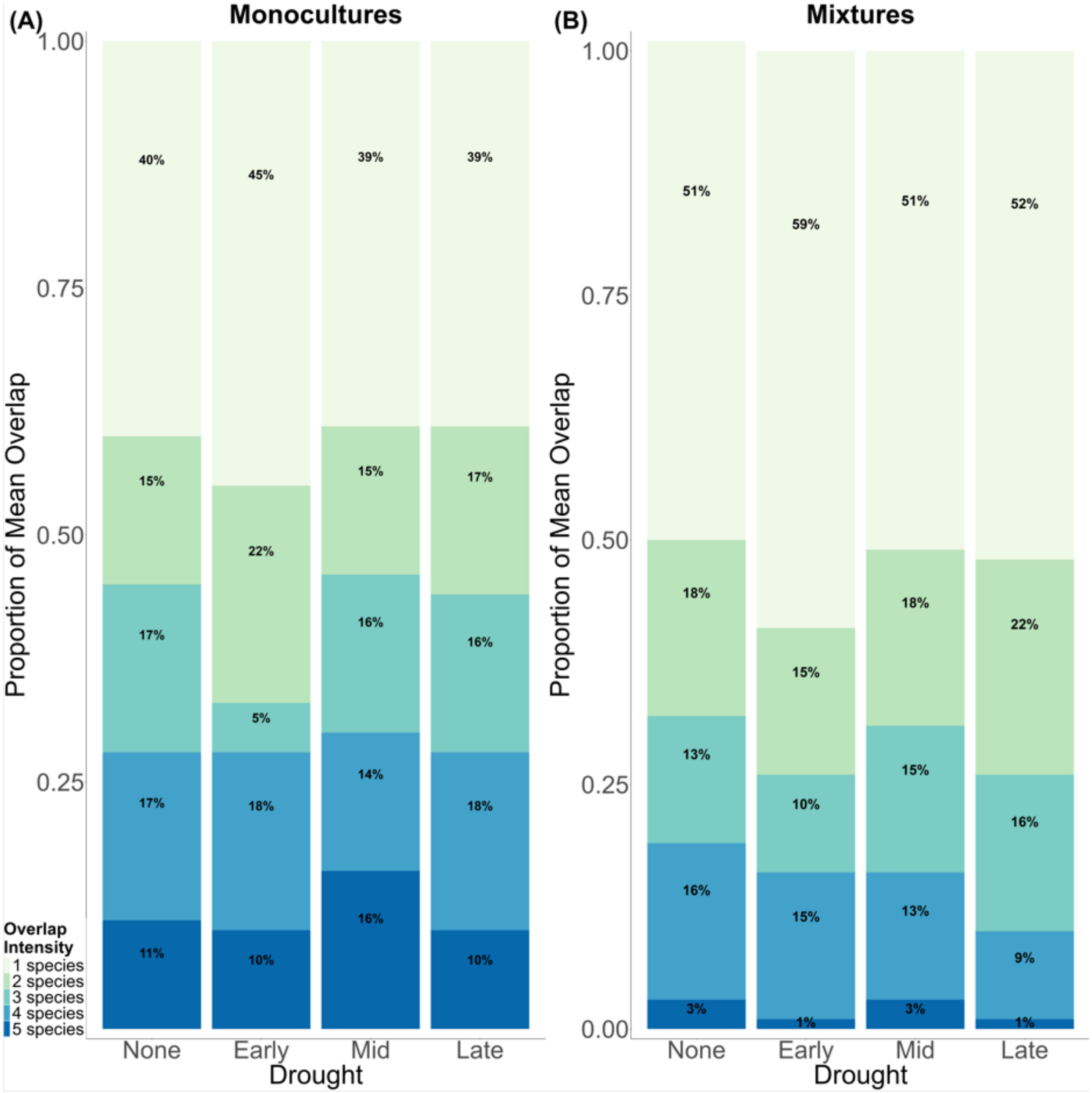


**Figure 4: The proportion of flowering weeks with different overlap intensities for each treatment.** (A) monocultures and (B) mixtures.

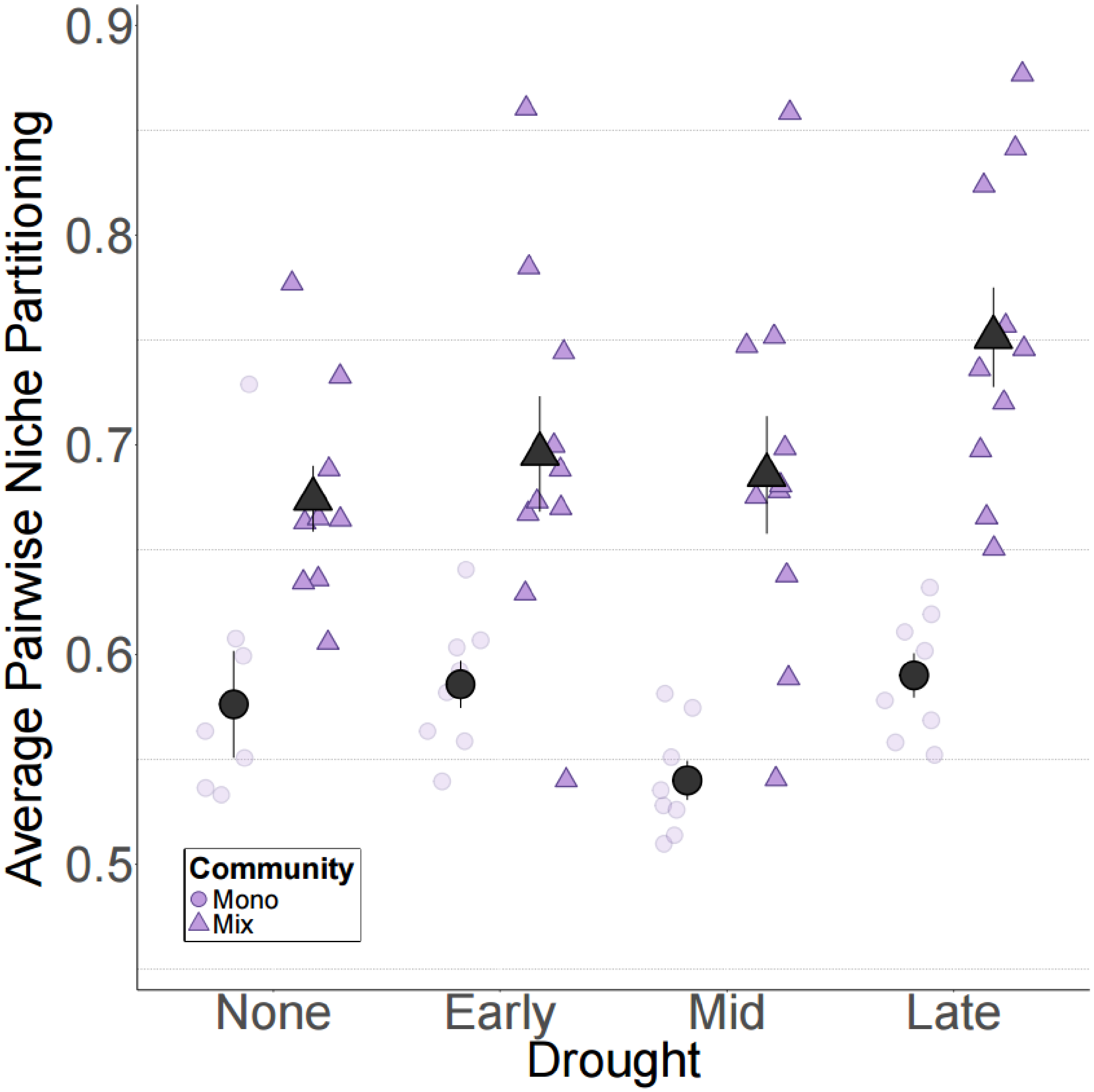


**Figure 5: The effect of treatments on the niche partitioning of whole communities (first year) as calculated by averaging the pairwise niche partitioning.** The y-axis is the averaged niche partitioning. The x-axis is the drought treatments. Round points are for the aggregated monocultures communities while triangular points are for the mixtures. The error bars are standard errors of means.

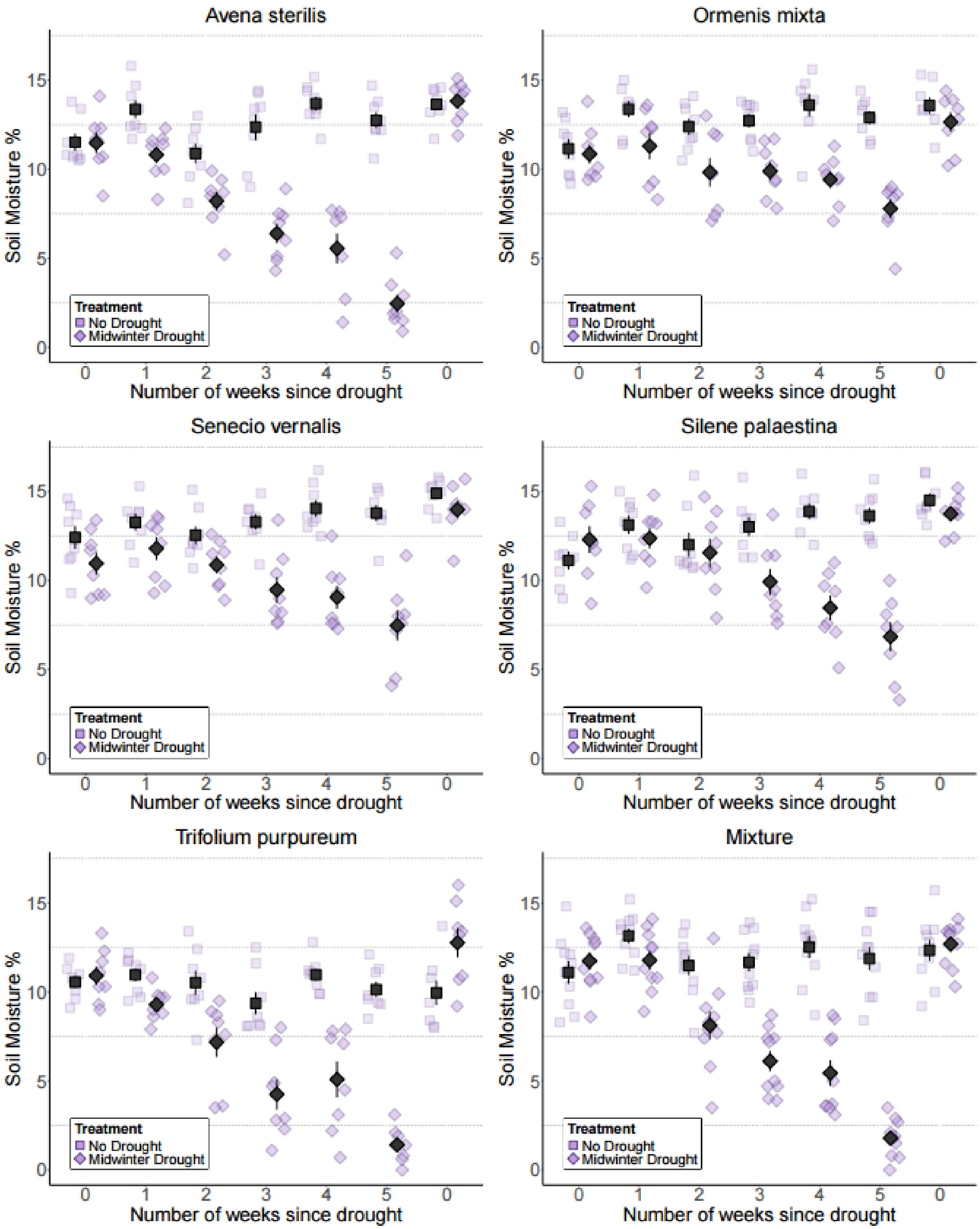


**Figure 6: The soil moisture content of monocultures and mixtures during midwinter drought and under control (first year).** The y-axis is the soil moisture percentage at a depth of 10cm. The x-axis is the number of weeks that have passed since the drought started (0 means the soil was sampled on a week with irrigation). Each treatment has its shape. The error bars are standard errors of means.

**Table 1:** Linear regression results for flowering duration across species. Models were fit to each species separately. Units are the number of weeks between the start and end of flowering. The parentheses show the SE and p-value for each predictor.

| Term | ***Avena sterilis*** | ***Ormenis mixta*** | ***Senecio vernalis*** | ***Silene palaestina*** | ***Trifolium purpureum*** |
|---|---|---|---|---|---|
| (Intercept) | 16.5*** (0.476, p<0.001) | 17.125*** (1.059, p<0.001) | 21.5*** (0.831, p<0.001) | 17.375*** (0.975, p<0.001) | 11.25*** (0.923, p<0.001) |
| Early Drought | -4.375*** (0.674, p<0.001) | -2.875. (1.498, p=0.06) | -3.625** (1.175, p=0.003) | -5** (1.379, p=0.001) | -3.125* (1.305, p=0.02) |
| Mid Drought | 0.25 (0.674, p=0.712) | 0.25 (1.498, p=0.868) | 2.75* (1.175, p=0.022) | -0.625 (1.379, p=0.652) | 1.125 (1.305, p=0.392) |
| Late Drought | -1.5* (0.674, p=0.029) | -0.375 (1.498, p=0.803) | 2.625* (1.175, p=0.029) | -1.375 (1.379, p=0.322) | -0.75 (1.305, p=0.568) |
| Mixture | -3*** (0.639, p<0.001) | -1.325 (1.421, p=0.355) | -4.5*** (1.115, p<0.001) | -7.875*** (1.308, p<0.001) | 1.15 (1.238, p=0.356) |
| Early Drought:Mixture | -1.225 (0.904, p=0.18) | -1.258 (2.035, p=0.538) | 1.025 (1.576, p=0.518) | 1 (1.85, p=0.591) | 0.625 (1.751, p=0.722) |
| Mid Drought:Mixture | -0.75 (0.904, p=0.41) | -0.75 (2.01, p=0.71) | -3.45* (1.576, p=0.032) | -0.775 (1.85, p=0.677) | 0.075 (1.751, p=0.966) |
| Late Drought:Mixture | 0.4 (0.904, p=0.66) | -1.125 (2.01, p=0.578) | -4.425** (1.576, p=0.007) | -1.458 (1.873, p=0.439) | -0.35 (1.751, p=0.842) |
| R-squared | 0.815 | 0.278 | 0.727 | 0.742 | 0.292 |
| No. of observations | 72 | 71 | 72 | 71 | 72 |

**Table 2:** Linear regression results for flowering start across species. Models were fit to each species separately. Units are the number of weeks since the first irrigation. The parentheses show the SE and p-value for each predictor.

| Term | ***Avena sterilis*** | ***Ormenis mixta*** | ***Senecio vernalis*** | ***Silene palaestina*** | ***Trifolium purpureum*** |
|---|---|---|---|---|---|
| (Intercept) | 10.625*** (0.34, p<0.001) | 22*** (0.623, p<0.001) | 4.75*** (0.144, p<0.001) | 13.625*** (0.761, p<0.001) | 17.375*** (0.549, p<0.001) |
| Early Drought | 5.625*** (0.48, p<0.001) | 2* (0.88, p=0.027) | 5.25*** (0.204, p<0.001) | 5.75*** (1.076, p<0.001) | 2.75** (0.777, p=0.001) |
| Mid Drought | 0.375 (0.48, p=0.438) | 0.125 (0.88, p=0.888) | -0.125 (0.204, p=0.542) | 0.625 (1.076, p=0.564) | 0.875 (0.777, p=0.264) |
| Late Drought | -0.25 (0.48, p=0.604) | -0.375 (0.88, p=0.672) | -0.125 (0.204, p=0.542) | 0.75 (1.076, p=0.489) | 0.5 (0.777, p=0.522) |
| Mixture | 2.475*** (0.456, p<0.001) | 0.5 (0.835, p=0.552) | -0.15 (0.193, p=0.441) | 3.775*** (1.021, p<0.001) | -0.075 (0.737, p=0.919) |
| Early Drought:Mixture | 0.475 (0.644, p=0.464) | 0.5 (1.196, p=0.677) | 0.15 (0.274, p=0.585) | -1.95 (1.444, p=0.182) | 0.15 (1.042, p=0.886) |
| Mid Drought:Mixture | 0.625 (0.644, p=0.336) | -0.125 (1.181, p=0.916) | 0.225 (0.274, p=0.414) | -0.025 (1.444, p=0.986) | -1.375 (1.042, p=0.192) |
| Late Drought:Mixture | 0.55 (0.644, p=0.397) | 0.075 (1.181, p=0.95) | 0.625* (0.274, p=0.026) | -1.039 (1.462, p=0.48) | -0.8 (1.042, p=0.445) |
| R-squared | 0.908 | 0.291 | 0.972 | 0.601 | 0.431 |
| No. of observations | 72 | 71 | 72 | 71 | 72 |

**Table 3:** Linear regression results for flowering end across species. Models were fit to each species separately. Units are the number of weeks since the first irrigation. The parentheses show the SE and p-value for each predictor.

| Term | ***Avena sterilis*** | ***Ormenis mixta*** | ***Senecio vernalis*** | ***Silene palaestina*** | ***Trifolium purpureum*** |
|---|---|---|---|---|---|
| (Intercept) | 26.125*** (0.385, p<0.001) | 38.125*** (0.749, p<0.001) | 25.25*** (0.872, p<0.001) | 30*** (0.554, p<0.001) | 27.625*** (0.764, p<0.001) |
| Early Drought | 1.25* (0.545, p=0.025) | -0.875 (1.059, p=0.412) | 1.625 (1.234, p=0.192) | 0.75 (0.784, p=0.342) | -0.375 (1.08, p=0.73) |
| Mid Drought | 0.625 (0.545, p=0.255) | 0.375 (1.059, p=0.724) | 2.625* (1.234, p=0.037) | 0 (0.784, p=1) | 2. (1.08, p=0.069) |
| Late Drought | -1.75** (0.545, p=0.002) | -0.75 (1.059, p=0.481) | 2.5* (1.234, p=0.047) | -0.625 (0.784, p=0.428) | -0.25 (1.08, p=0.818) |
| Mixture | -0.525 (0.517, p=0.313) | -0.825 (1.004, p=0.415) | -4.65*** (1.17, p<0.001) | -4.1*** (0.744, p<0.001) | 1.075 (1.025, p=0.298) |
| Early Drought:Mixture | -0.75 (0.731, p=0.309) | -0.758 (1.438, p=0.6) | 1.175 (1.655, p=0.48) | -0.95 (1.052, p=0.37) | 0.775 (1.449, p=0.595) |
| Mid Drought:Mixture | -0.125 (0.731, p=0.865) | -0.875 (1.42, p=0.54) | -3.225. (1.655, p=0.056) | -0.8 (1.052, p=0.45) | -1.3 (1.449, p=0.373) |
| Late Drought:Mixture | 0.95 (0.731, p=0.198) | -1.05 (1.42, p=0.463) | -3.8* (1.655, p=0.025) | -2.497* (1.065, p=0.022) | -1.15 (1.449, p=0.43) |
| R-squared | 0.424 | 0.198 | 0.671 | 0.772 | 0.175 |
| No. of observations | 72 | 71 | 72 | 71 | 72 |

**Table 4:** Linear regression results for niche partitioning across treatments. This table summarizes 100 regression analyses — each one ran with the same 40 mixtures but different sets of 32 independent simulated communities ("Simulated"). The presented predictors and p-values are calculated based on these 100 models (see details in methods). Units are the niche partitioning value (0–1).

| Term | ***Whole communities*** |
|---|---|
| (Intercept) | 0.747*** ($p<0.001$) |
| Early Drought | 0.042* ($p=0.033$) |
| Mid Drought | 0.007 ($p=0.737$) |
| Late Drought | 0.043* ($p=0.029$) |
| Simulated | -0.104*** ($p<0.001$) |
| Early Drought:Simulated | -0.002 ($p=0.941$) |
| Mid Drought:Simulated | -0.028 ($p=0.343$) |
| Late Drought:Simulated | -0.043 ($p=0.138$) |

**Table 5:** Linear regression results for pairwise average niche partitioning across treatments. The model was fit for the 40 mixtures and one set of 32 independent communities. Units are the mean calculated niche partitioning value (0–1). The parentheses show the SE and p-value for each predictor.

| Term | ***Pairwise*** |
|---|---|
| (Intercept) | 0.674*** (0.02, p<0.001) |
| Early Drought | 0.021 (0.029, p=0.462) |
| Mid Drought | 0.011 (0.029, p=0.694) |
| Late Drought | 0.077* (0.029, p=0.01) |
| synthetic | -0.098** (0.031, p=0.002) |
| Early Drought:synthetic | -0.012 (0.043, p=0.785) |
| Mid Drought:synthetic | -0.048 (0.043, p=0.275) |
| Late Drought:synthetic | -0.063 (0.043, p=0.149) |
| R-squared | 0.562 |
| No. of observations | 72 |
